# Limitations of proposed signatures of Bayesian confidence

**DOI:** 10.1101/218222

**Authors:** William T. Adler, Wei Ji Ma

## Abstract

The Bayesian model of confidence posits that confidence is the observer’s posterior probability that the decision is correct. It has been proposed that researchers can gain evidence in favor of the Bayesian model by deriving qualitative signatures of Bayesian confidence, i.e., patterns that one would expect to see if an observer was Bayesian, and looking for those signatures in human or animal data. We examine two proposed qualitative signatures, showing that their derivations contain hidden assumptions that limit their applicability, and that they are neither necessary nor sufficient conditions for Bayesian confidence. One signature is an average confidence of 0.75 for trials with neutral evidence. This signature only holds when class-conditioned stimulus distributions do not overlap and internal noise is very low. Another signature is that, as stimulus magnitude increases, confidence increases on correct trials but decreases on incorrect trials. This signature is also dependent on stimulus distribution type. There is an alternative form of this signature that has been applied in the literature; we find no indication that it is expected under Bayesian confidence, which resolves an ostensible discrepancy. We conclude that, to determine the nature of the computations underlying confidence reports, there may be no shortcut to quantitative model comparison.

## 1 Introduction

Humans possess a sense of confidence about decisions they make, and asking human subjects for their decision confidence has been a common psychophysical method for over a century^33^. But despite the long history of confidence reports, it is still unknown how the brain computes confidence reports from sensory evidence. The leading proposal has been that confidence reports are a function of the observer’s posterior probability of being correct ^10,14,17,28,34^, a hypothesis we call the Bayesian confidence hypothesis (BCH)^1^.

In recent years, some researchers have tested the BCH by formally comparing Bayesian confidence models to other models^1,3^. Although this is the most thorough method to test the BCH, it can be painstaking in practice. To avoid this approach, one could instead try to describe qualitative patterns that should theoretically emerge from Bayesian confidence and then look for those patterns in real data. Partly following this motivation, Hangya et al.^14^ propose signatures of the BCH, some of which have been observed in behavior^18,22,37^ and in neural activity^18,21^.

These signatures are not unique to the Bayesian model; instead, they are expected under a number of other models^17^. This may be considered an advantage for a confidence researcher who is not interested in the precise algorithmic underpinnings of confidence. A researcher may observe these signatures in behavior, reasonably conclude that she has evidence that the organism is computing some form of confidence, and probe more deeply into, for instance, neural activity^18^. In this manuscript, however, we consider the researcher concerned with understanding the algorithm used by an organism to compute confidence. For such a researcher, the fact that these signatures emerge from multiple models poses a problem: These signatures are not sufficient conditions for any particular model of confidence, including the Bayesian model. In other words, observation of these signatures does not constitute strong evidence in favor of any particular model. Because of this insufficiency, we view with skepticism any research that uses observation of these signatures as the basis for a claim that an organism uses a Bayesian form^29^, “statistical” form^37^, or any other specific form of confidence.

Although they do not claim that the signatures are sufficient conditions, Hangya et al. do claim that the signatures are necessary conditions for the BCH, i.e., that if confidence is Bayesian, these patterns will be present in behavior. If the signatures are necessary but not sufficient conditions for the BCH, observation of a single signature does not imply that the BCH is true; instead, one would need to observe several signatures in order to gain confidence in the nature of confidence.^I^ However, we show that two of these signatures are not necessary conditions, reducing the overall value of the qualitative signature method for testing the BCH.

One signature is a mean confidence (i.e., the observer’s estimated probability of being correct) of 0.75 for trials with neutral evidence. We show that, under the Bayesian model, this signature will only be observed when noise is very low and stimulus distributions do not overlap.

Another signature is that, as stimulus magnitude increases, mean confidence increases on correct trials but decreases on incorrect trials. Here, we show that under the Bayesian model, this signature breaks down when noise is low and stimulus distributions are Gaussian. We also explain and resolve a recent discrepancy in the literature that is related to an alternative formulation of this signature^29^.

## 2 Binary categorization task

We restrict ourselves to the following, widely used, family of binary perceptual categorization tasks^13^. On each trial, a category *C* ∈ {−1,1} is randomly drawn with equal probability. Each category corresponds to a stimulus distribution *p*(*s* | *C*), where *s* may specify the value of many possible kinds of stimuli (e.g., an odor mixture^18^, the net motion energy of a random dot kinematogram^19,30^, the orientation of a Gabor^1,7,35^, or the mean orientation of a series of Gabors^29^). The stimulus distributions are mirrored across *s* = 0, i.e., *p*(*s* | *C* = −1) = *p*(−*s* | *C* = 1). We assume that the observer has full knowledge of these distributions. A stimulus *s* is drawn from the chosen stimulus distribution and presented to the observer. The observer does not have direct access to the value of *s*; instead, they take a noisy measurement *x*, drawn from the distribution 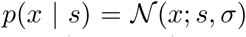 distribution over *x* with mean *s* and standard deviation *σ* (Figure 1).

**Figure 1:**
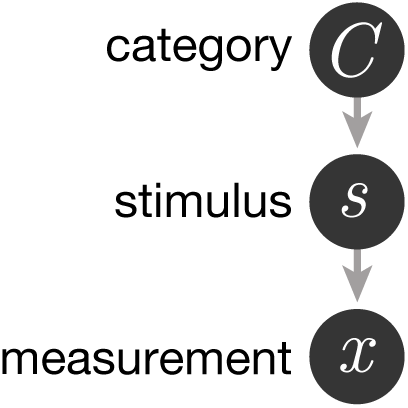
Generative model of the task.

If the observer’s choice behavior is Bayes-optimal (i.e., minimizes expected loss which, in a task where each category has equal reward, is equivalent to maximizing accuracy), they compute the posterior probability of each category by marginalizing over all possible values of *s*: 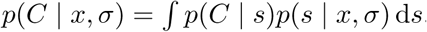. They then make a category choice 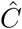 by choosing the category with the highest posterior: 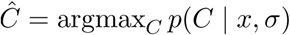. For mirrored stimulus distributions, that amounts to choosing 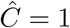 when *x* > 0, and 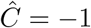 otherwise.

Furthermore, if the observer’s confidence behavior is Bayesian, then it will be some function of the posterior probability of the chosen category. This probability is 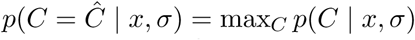. Because it is a deterministic function of *x* and *σ*, we will refer to it as conf(*x*,*σ*).^II^ See Appendix A for derivations of conf(*x*,*σ*) for all stimulus distribution types used in this paper.

## 3 0.75 signature: Mean Bayesian confidence is 0.75 for neutral evidence trials

Hangya et al.^14^ propose a signature concerning neutral evidence trials, those in which the stimulus *s* is equal to 0 (i.e., there is equal evidence for each category), and observer performance is at chance. Bayesian confidence on each individual trial will always be at least 0.5 (assuming that measurement noise is nonzero). One can intuitively understand why this is: in binary categorization, if the posterior probability of one option is less than 0.5, the observer makes the other choice, which has a posterior probability above 0.5. Therefore, all trials have confidence of at least 0.5, and mean confidence at any value of *s* is also greater than 0.5. Hangya et al. go beyond these results and provide a proof that, under some assumptions, mean Bayesian confidence on neutral evidence trials is *exactly* 0.75. We refer to this prediction as the 0.75 signature, and we show that it is not always expected under a normative Bayesian model.

### 3.1 The 0.75 signature is not a necessary condition for Bayesian confidence

To determine the conditions under which the 0.75 signature is expected under the Bayesian model, we used Monte Carlo simulation with the following procedure. For a range of measurement noise levels *σ*, we drew measurements *x* from 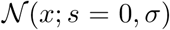. Using the function conf(*x*, *σ*) that the observer would use if they believed stimuli were being drawn from category-conditioned stimulus distributions *p*(*s* | *C*) (rather than all *s* being zero), we computed Bayesian confidence for each measurement. We then took the mean confidence, equal to 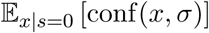.

The 0.75 signature only holds if the s.d. of the noise is very low relative to the range of the stimulus distribution. Additionally, the observer must believe that the category-conditioned stimulus distributions are non-overlapping (Figure 2a, dotted line). If the observer believes that the category-conditioned stimulus distributions overlap by even a small amount, mean confidence on neutral evidence trials drops to 0.5. Therefore, in an experiment with overlapping stimulus distributions, one should not expect an optimal observer to produce the 0.75 signature. In experiments with non-overlapping distributions, an observer’s false belief about the distributions might also cause them to not produce the 0.75 signature. We use the example of overlapping uniform stimulus distributions (Figure 2a, solid lines) to demonstrate the fragility of this signature, although such distributions are not common in the literature. Overlapping Gaussian stimulus distributions (Figure 2b), however, are relatively common in the perceptual categorization literature^1,4,13,31,35^ and arguably more naturalistic^26^. Because the 0.75 signature requires both low measurement noise and non-overlapping stimulus distributions, mean 0.75 confidence at neutral evidence trials is not a necessary condition for Bayesian confidence.

Additionally, the 0.75 signature is only relevant in experiments where subjects are specifically asked to report confidence in the form of a perceived probability of being correct (or are incentivized to do so through a scoring rule^6,12,27^, although in this case it has been argued^1,25^ that any Bayesian behavior might simply be a learned mapping). In other words, in an experiment where subjects are asked to report confidence on a 1 through 5 scale, a mean confidence of 3 only corresponds to 0.75 if one makes the a priori assumption that there is a linear mapping between rating and perceived probability of being correct^37^.

**Figure 2:**
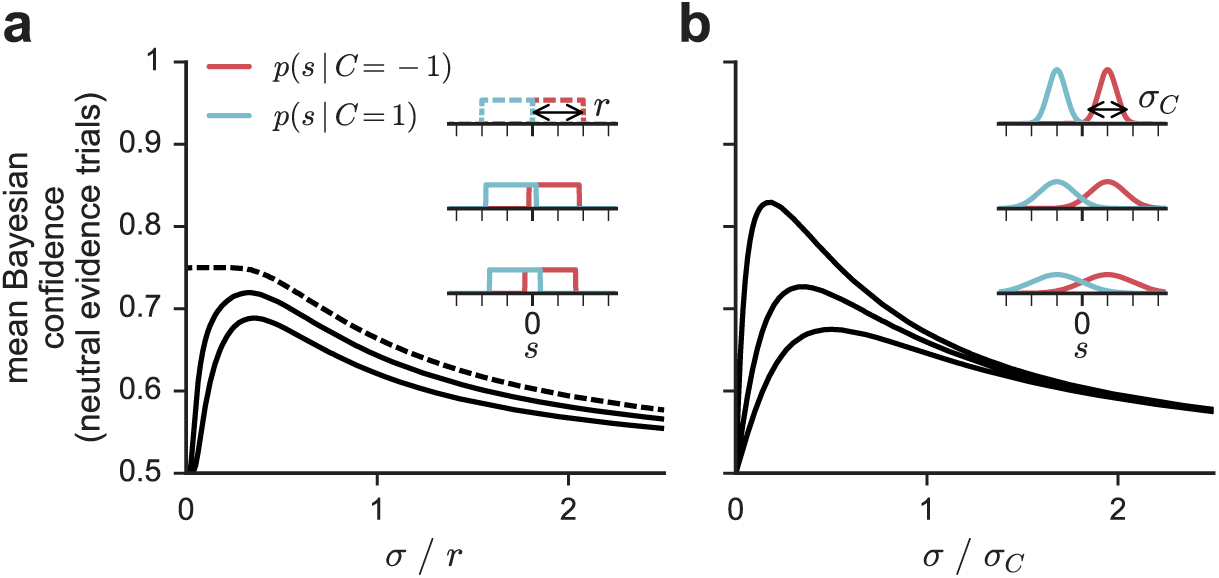
The 0.75 signature is not a necessary condition for Bayesian confidence. The y-axis indicates mean Bayesian confidence on trials for which *s* = 0. Each inset corresponds to a line, in the same top-to-bottom order. Dotted and solid lines indicate, respectively, non-overlapping and overlapping categories. For each value of *σ*, 50,000 trials were simulated. (**a**) Trials were simulated using uniform stimulus distributions defined by 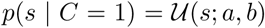, with *b* − *a* = *r* = 2. When the stimulus categories are non-overlapping (i.e., with *a* = 0 and *b* = 2, top inset), the 0.75 signature can be observed at zero measurement noise (dotted black line). However, mean Bayesian confidence decreases as a function of measurement noise. Additionally, when the distributions overlap slightly (bottom two insets), the 0.75 signature will not be observed (solid black lines). (**b**) Moreover, when the stimulus categories are Gaussian distributions defined by 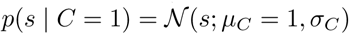, the 0.75 signature will not be observed at any *σ*_*C*_ or measurement noise level *σ*. One can intuitively understand why mean confidence is 0.5 for overlapping categories at very low measurement noise and increases with measurement noise. At very low measurement noise, the observer makes measurements that are very close to zero, which the observer “knows” are associated with a low probability of being correct. However, as noise increases, the observer starts to make measurements that have higher magnitude, leading the observer to believe that they have a higher probability of being correct.

#### 3.1.1 Relevant assumptions in Hangya et al

Hangya et al. describe an assumption that is critical for the 0.75 signature: each category-conditioned stimulus distribution is a continuous uniform distribution. However, the 0.75 signature depends on two additional assumptions that they make implicitly.

Their proof depends on confidence for one category being equal to *p*(*s* > 0 | *x*, *σ*) (p. 1852). This equality further depends on their implicit assumption both of non-overlapping categories and of negligible measurement noise; these assumptions are equivalent to only considering the leftmost point of the solid line in Figure 2a. To understand why, we derive their definition of confidence as *p*(*s* > 0 | *x*, *σ*).

Without loss of generality, we look at trials with choice 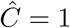. First, Hangya et al. make the assumption that the categories are non-overlapping uniforms (i.e., 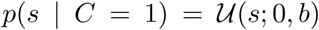, which denotes a continuous uniform distribution over *s* between 0 and *b*). This allows them to write (Appendix A):

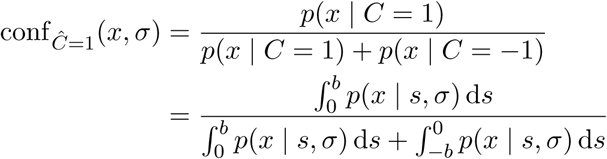

Second, they make the assumption that *b* is very large relative to measurement noise *σ*. This allows them to write:

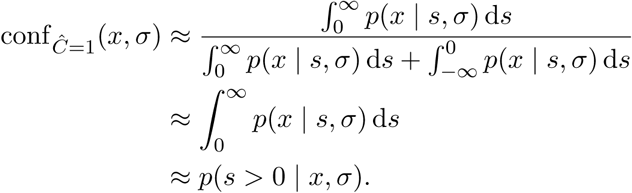

If the stimulus distributions overlap by even a small amount or if measurement noise is non-negligible, confidence cannot be written as *p*(*s* > 0 | *x*, *σ*), and the proof of the 0.75 signature breaks down.

### 3.2 The 0.75 signature is not a sufficient condition for Bayesian confidence

We have shown that the 0.75 signature is not a necessary condition for Bayesian confidence, but is it a sufficient condition? It is possible to show that a signature is a sufficient condition if it is not possible to observe it under any other model. However, one could put forward a trivial model that always produces exactly midrange confidence on each trial, regardless of the measurement. Therefore, the 0.75 signature is not a sufficient condition.

## 4 Divergence signature #1: As stimulus magnitude increases, mean confidence increases on correct trials but decreases on incorrect trials

Hangya et al.^14^ propose the following pattern as a signature of Bayesian confidence: On correctly categorized trials, mean confidence is an increasing function of stimulus magnitude (here, |*s*|), but on incorrect trials, it is a decreasing function (Figure 3a). We refer to this pattern as the divergence signature.^III^ The signature is present in Bayesian confidence when category-conditioned stimulus distributions are uniform, in both high- and low-noise regimes (Figure 3a,b). The intuition for why this pattern may occur is as follows. On correct trials, as stimulus magnitude increases, the mean magnitude of the measurement *x* increases. Because measurement magnitude is monotonically related to Bayesian confidence, this increases mean confidence. However, on incorrect trials (in which *x* and *s* have opposite signs), the mean magnitude of the measurement decreases (Figure 5a), which in turn decreases mean confidence (Figure 5b,c).

The divergence signature has been observed in some behavioral experiments^18,21,22,37^. However, we demonstrate that, as with the 0.75 signature the divergence signature is not always expected under a normative Bayesian model.^IV^ Therefore, the appearance of the signature in these papers should not be taken to mean that it should be generally expected.

### 4.1 Divergence signature #1 is not a necessary condition for Bayesian confidence

To determine the conditions under which the divergence signature is expected under the Bayesian model, we used Monte Carlo simulation with the following procedure. We generated stimuli *s*, drawn with equal probability from stimulus distributions *p*(*s* | *C* = −1) and *p*(*s* | *C* = 1). We generated noisy measurements *x* from these stimuli, using measurement noise levels *σ*. We generated observer choices from these measurements, using the optimal decision rule 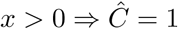, and we computed Bayesian confidence for every trial.

When stimulus distributions are Gaussian and measurement noise is low relative to stimulus distribution width, the divergence signature is not expected (Figure 3c,d). To understand why this is, imagine an optimal observer with zero measurement noise. In tasks with overlapping categories, even this observer cannot achieve perfect performance; for a given category with a positive mean, there are stimuli that have a negative value, resulting in an incorrect choice. For such stimuli, confidence *increases* with stimulus magnitude. At relatively low noise levels, these stimuli represent the majority of all incorrect trials for the category (Figure 3e). This effect causes the divergence signature to disappear when averaging over trials drawn from both categories. Because of this, the divergence signature is not a necessary condition for Bayesian confidence. Note that an experimenter could avoid this issue by plotting confidence as a function of signed stimulus value *s* and by not averaging over both categories, which would produce plots such as Figure 3e.

#### 4.1.1 Relevant assumption in Hangya et al

We have shown that the applicability of the divergence signature may be limited to particular cases. By contrast, the proof in Hangya et al. suggests that it is quite general. We can resolve this paradox by making explicit the assumptions hidden in the proof. They assume that, “for incorrect choices…with increasing evidence discriminability, the relative frequency of low-confidence percepts increases while the relative frequency of high-confidence percepts decreases” (p. 1847).^V^ This assumption is violated in the case of overlapping Gaussian stimulus distributions: for some incorrect choices (branch 4 of Figure 3e), as *s* becomes more discriminable (i.e., very negative), the frequency of *high*-confidence reports increases. At low levels of measurement noise, this causes the divergence signature to disappear.

**Figure 3:**
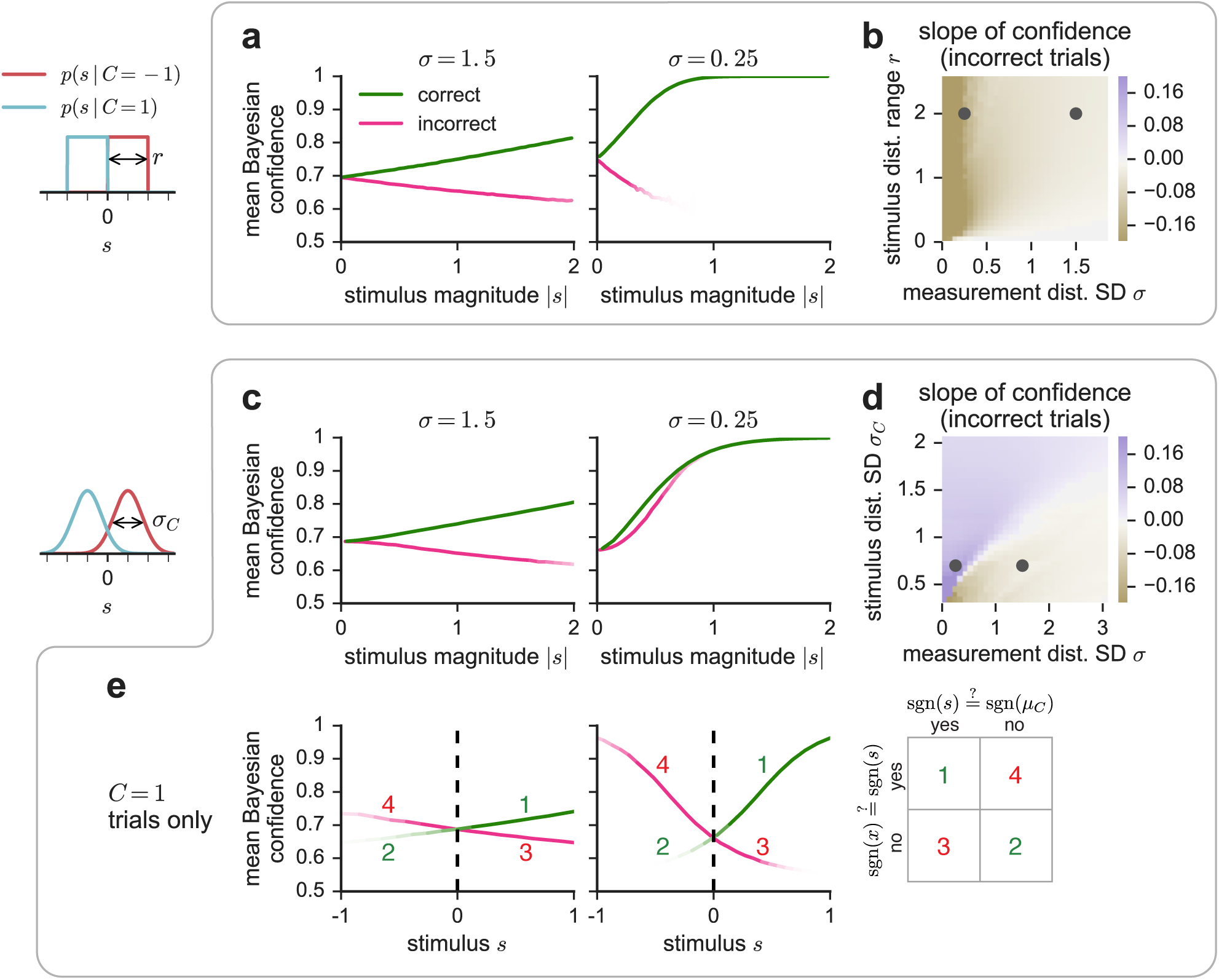
The divergence signature is not a necessary condition for Bayesian confidence. For two stimulus distribution types, we simulated 2 million trials. (**a**) With uniform stimulus distributions defined by 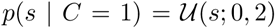, the divergence signature is predicted under both high- and low-noise regimes. The fadedness of the line indicates conditions for which there are few trials. (**b**) Heatmap indicates the slope of the pink lines in **a**. At all values of *σ* and distribution range, the slope is negative. Slopes were obtained by generating binned mean confidence values as in **a** and fitting a line to those values. Black markers indicate the parameters used in **a**, with left dot corresponding to right plot and conversely. (**c**) With Gaussian stimulus distributions defined by 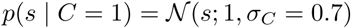, the divergence signature appears only when measurement noise is high, i.e., when *σ* ≲ 0.6. (**d**) As in **b** but for Gaussian distributions with means of ±1. Under some values of *σ* and *σ*_*C*_, the slope is positive, indicating that the divergence signature is not a necessary condition for Bayesian confidence. (**e**) Visual explanation for why, under Gaussian stimulus distributions, the divergence signature appears only at relatively high *σ* values. Plots represent the same data as in **c**, but over *s* instead of |*s*|. For clarity, we only use trials drawn from category *C* = 1; the argument is unaffected. Incorrect trials fall into two categories: on trials in which *s* is positive but *x* is negative due to noise, confidence goes down as |*s*| increases (branch 3); on trials in which *s* and *x* are both negative, confidence increases with |*s*| (branch 4). At high levels of noise, branch 3 has more trials than branch 4, and dominates the averaging that occurs when plotting trials from both categories over |*s*|. At low levels of noise, branch 4 instead dominates, and the divergence signature disappears. Note that, for non-overlapping distributions (e.g., those in **a**,**b**), there are no trials in which *s* has a different sign than the stimulus distribution mean, so branches 2 and 4 do not exist, and the divergence signature is always present.

### 4.2 Divergence signature #1 is not a sufficient condition for Bayesian confidence

It has been previously noted that the signature is expected under a number of non-Bayesian models^11,16,17^. Here, we describe an additional non-Bayesian model, one in which confidence is a function only of |*x*|, the magnitude of the measurement.^VI^ In the general family of binary categorization tasks described in Section 2, the confidence of this model is monotonically related to the confidence of the Bayesian model conf(*x*,*σ*). Thus, when the divergence signature is predicted by the Bayesian model, it is also predicted by this measurement model. Therefore, the divergence signature is not a sufficient condition for Bayesian confidence.

## 5 Divergence signature #2: As measurement noise decreases, mean confidence increases on correct trials but decreases on incorrect trials

An alternative version of the divergence signature has emerged in the literature. Navajas et al.^29^ conduct an experiment in which they present, on each trial, a series of oriented Gabors with orientations pseudorandomly drawn from uniform distributions with different variances. They then ask subjects to judge whether the mean orientation is left or right of vertical and to provide a confidence report. They plot confidence as a function of correctness and orientation distribution variance, expecting that, if confidence were Bayesian, their data would look like Figure 3a. Contrary to their expectations, they observe no such divergence (Figure 4a). However, instead of plotting *stimulus magnitude* which produces divergence signature #1 (Section 4), they plot *measurement noise*^VII^ on the x-axis (Figure 4a), in effect proposing a divergence signature distinct from the one described in Section 4. We will refer to this as divergence signature #2: as measurement noise decreases, mean confidence increases on correct trials but decreases on incorrect trials. We find no evidence that this signature is expected under the Bayesian confidence model, resolving the seemingly unexpected result in Navajas et al.

### 5.1 Divergence signature #2 is not expected under Bayesian confidence

To determine whether divergence signature #2 is expected under the Bayesian model, we used Monte Carlo simulation with the following procedure. We generated stimuli with *s* = ±1, corresponding to *C* = ±1.^VIII^ For a range of measurement noise levels *σ*, we drew noisy measurements *x* from 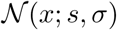. We generated observer choices from these measurements, using the optimal decision rule 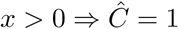. We computed Bayesian confidence for every trial.

As measurement noise decreases, mean confidence increases for both correct and incorrect trials (Figure 4b). This pattern also holds when the category-conditioned stimulus distributions are uniform or Gaussian, and if one plots a measure of stimulus distribution variance on the x-axis (either uniform distribution range *r* or Gaussian distribution s.d. *σ_C_*). This indicates that the signature is not expected under the BCH.

**Figure 4:**
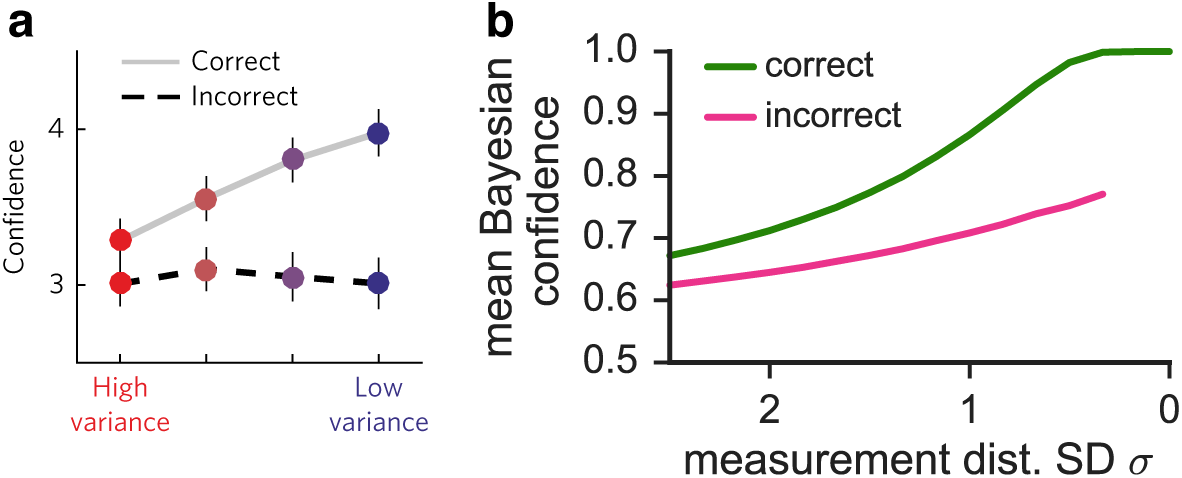
Divergence signature #2 is not present either in the Navajas et al. data or in the prediction of the Bayesian model. (**a**) Average confidence in a binary perceptual categorization task, reproduced with permission from Navajas et al. ^29^. (**b**) Mean Bayesian confidence as a function of measurement noise is not expected to show opposite trends when conditioned on correctness. At each value of *σ*, 50,000 stimuli were stimulated, with *s* = ±1.

#### 5.1.1 Related text in Hangya et al

It is quite understandable that Navajas et al. took measurement noise as their definition of evidence discriminability; Hangya et al. explicitly allow it in their description of the divergence signature: “any monotonically increasing function of expected outcome [i.e., accuracy]…can serve as evidence discriminability” (p. 1847). Measurement noise (or, in keeping strictly with Hangya et al.‘s definition, measurement precision) is indeed monotonically related to accuracy. However, the divergence signature requires an additional assumption: “for incorrect choices…with increasing evidence discriminability, the relative frequency of low-confidence percepts increases while the relative frequency of high-confidence percepts decreases” (p. 1847; see also, Section 4.1.1). Simulation shows that this assumption is violated when measurement noise is used as the definition of evidence discriminability.

#### 5.1.2 Why the intuition for divergence signature #1 does not predict divergence signature # 2

We have shown that, although d-vergence signature #1 is not completely general, it is expected under the Bayesian model in some cases (Figure 3a). By contrast, there is no indication that divergence signature #2 is ever expected. This may be surprising, because the intuition for divergence signature #1 might seem to apply equally to divergence signature #2. However, the effect of measurement noise on mean confidence is different than the effect of stimulus magnitude because measurement noise, unlike stimulus magnitude, affects the mapping from measurement to confidence on a single trial.

Mean Bayesian confidence is a function of two factors: confidence on a single trial and the probability of the corresponding measurement.

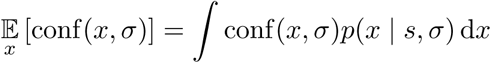

The intuition for divergence signature #1 is as follows: as stimulus magnitude |*s*| increases, the measurement distribution *p*(*x* | *s*, *σ*) shifts, and the mean measurement magnitude on incorrect trials decreases (Figure 5a). One might expect this intuition to also result in divergence signature #2, since the effect of decreased measurement noise *σ* on *p*(*x* | *s*, *σ*) also results in a decreased measurement magnitude on incorrect trials (Figure 5d). However, *σ* additionally affects conf(*x*,*σ*), the per-trial, deterministic mapping from measurement and noise level to Bayesian confidence (Figure 5e), whereas stimulus magnitude does not (Figure 5b). Therefore, when *σ* is variable, the resulting effect on the measurement distribution is insufficient for describing the pattern of mean confidence on incorrect trials, requiring simulation. We simulated experiments as described in Section 4.1, and demonstrate why stimulus magnitude and measurement noise have different effects on mean confidence on incorrect trials (Figure 5).

**Figure 5:**
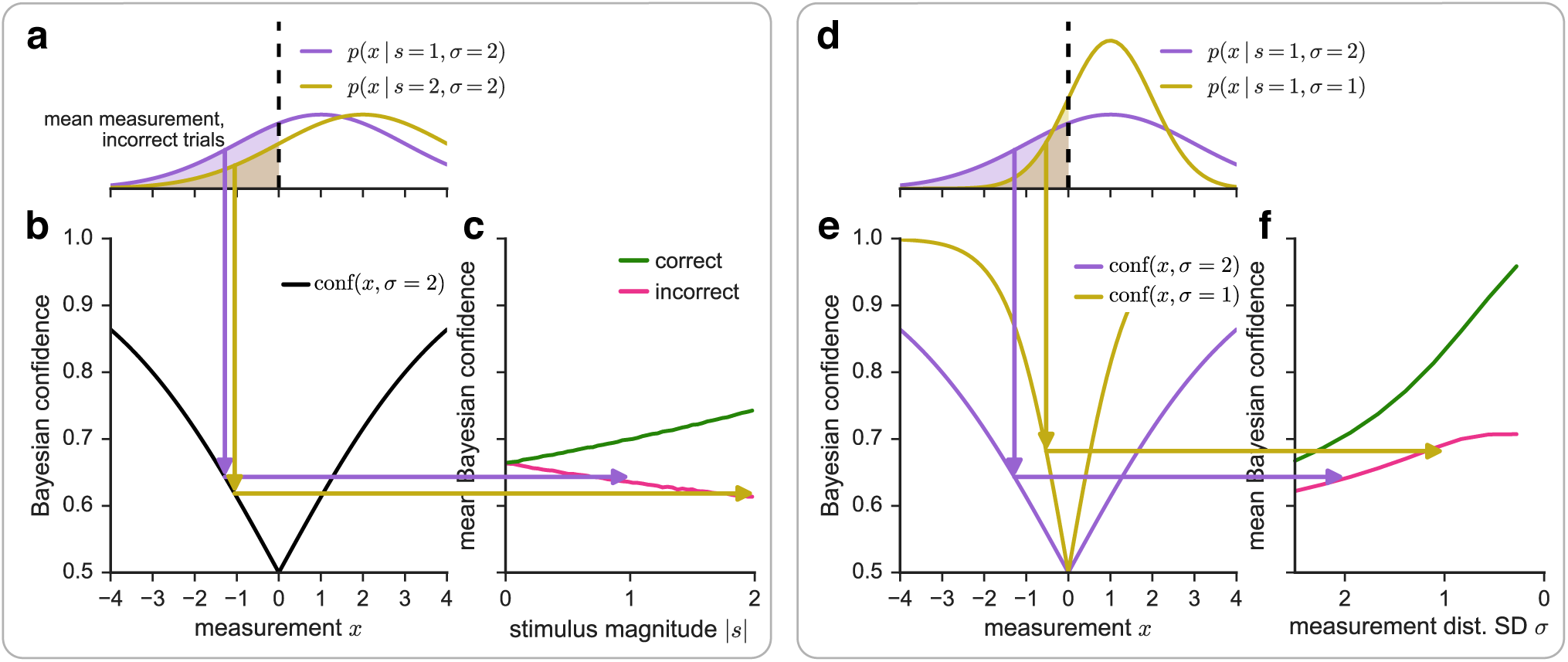
Explanation for why divergence signature #1 is sometimes expected, but why divergence signature #2 might not ever be expected. Although increased stimulus magnitude and decreased measurement noise both cause the mean measurement magnitude to decrease on incorrect trials, they have different effects on mean confidence. At each value of *σ* 2 million stimuli were simulated, using uniform stimulus distributions defined by 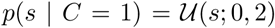 (the case of Figure 3a). (**a**) As described previously^8,14,18^, an increase in stimulus magnitude causes the mean measurement magnitude to decrease on incorrect trials. (**b**) Measurements are mapped onto confidence values using the deterministic function conf(*x*, *σ*), which is equivalent to the posterior probability that the choice is correct (Section 2). (**c**) This mapping results in divergence signature #1, a decrease in mean confidence on incorrect trials. Arrows do not align precisely with the simulated mean, because the confidence of the mean measurement is not exactly equal to the mean confidence. (**d**) A decrease in measurement noise also causes the mean measurement magnitude to decrease on incorrect trials. (**e**) Because the mapping from measurement to confidence conf(*x*,*σ*) is dependent on *σ*, measurements from the less noisy distribution have higher confidence. (**f**) Because the confidence mapping is dependent on *σ*, divergence signature #2 is not necessarily expected under Bayesian confidence.

### 5.2 Use of divergence signature #2 in Navajas et al

Navajas et al. motivate their findings by first building a *non-Bayesian* model of confidence^IX^ that does predict divergence signature #2, i.e., that, as measurement noise decreases, mean confidence decreases on incorrect trials. They then fail to observe the signature in their averaged data (Figure 4a), observing instead that confidence is constant on incorrect trials. Some subjects (e.g., subject 16 in their Figure 3), however, do show the signature. This leaves them with a puzzle—what model can describe the data? To answer this, they modify their model to incorporate Fisher information, which increases as measurement noise decreases. This post-hoc model is able to “bend” the confidence curve upward as measurement noise decreases, producing curves that more closely resemble their data.

The main shortcoming of this argument is that a Bayesian model of confidence would not actually predict divergence signature #2, as we have shown above. Indeed, their averaged data more closely resembles the prediction of the Bayesian model (Figure 4b) than that of their non-Bayesian model without Fisher information (their Figure 2b). Therefore, the absence of the signature in their averaged data does not suggest anything beyond a Bayesian model; it is possible that the Bayesian model would provide a good fit to most of their subjects. If the model provided a poor fit to subjects that do show divergence signature #2, a post-hoc model would have to incorporate some other mechanism that could “bend” the confidence curve *downward* which would not be Fisher information.

## 6 Other signatures

A third signature in Hangya et al.^14^ that we do not discuss here (that confidence equals accuracy), is like the 0.75 signature in that it either requires explicit reports of perceived probability of being correct, or the experimenter to choose a mapping between rating and perceived probability of being correct (Section 3.1). For any monotonic relationship between accuracy and confidence, it is likely that there is some mapping that equates the two, in which case the signature would not be a sufficient condition for the BCH.

A fourth signature (that confidence allows a better prediction of accuracy than stimulus magnitude alone) is, like divergence signature #1, also predicted by the measurement model (Section 4.2) and is therefore also not a sufficient condition for the BCH.

## 7 Discussion

We have demonstrated that, even in the relatively restricted class of binary categorization tasks that we consider here (Section 2), some signatures are neither necessary nor sufficient conditions for the BCH. Specifically, the 0.75 signature is only expected under non-overlapping stimulus distributions. Additionally, despite claims that divergence signature #1 is “robust to different stimulus distributions,” ^17^ it is only expected under non-overlapping stimulus distributions or under Gaussian stimulus distributions with high measurement noise. Because of their non-generality, these signatures are therefore not necessary conditions of Bayesian confidence. Furthermore, they may be observed under non-Bayesian models, indicating that they are also not sufficient conditions^11,16^.

A discrepancy in the literature^29^ has emerged through the confusion of divergence signature #1 with a second form, in which stimulus magnitude is replaced with measurement noise.^X^ We have shown that, while divergence signature #1 holds in some cases, there is no evidence that the second form is ever expected under the BCH, which resolves this discrepancy.

Some of our critique of the signatures has focused on the implicit assumption that experiments use non-overlapping stimulus distributions. One could object to our critique by questioning the relevance of overlapping stimulus distributions, given that non-overlapping stimulus distributions are the norm in the confidence literature^2,17,18,37^. But although overlapping categories are only just beginning to be used to study confidence^1,7^, such categories have a long history in the perceptual categorization literature^4,13,15,23,24,35,36^. It has been argued that overlapping Gaussian stimulus distributions have several properties that make them more naturalistic than non-overlapping distributions^26^. The property most relevant here is that with overlapping categories, perfect performance is impossible, even with zero measurement noise. With overlapping categories, as in real life, identical stimuli may belong to multiple categories. Imagine a coffee drinker pouring salt rather than sugar into her drink, a child reaching for his parent’s glass of whiskey instead of his glass of apple juice, or a doctor classifying a malignant tumor as benign^5^. In all three examples, stimuli from opposing categories may be visually identical, even under zero measurement noise. For more naturalistic experiments with overlapping categories, qualitative signatures will be unusable if their derivations assume non-overlapping categories.

Given our demonstration that proposed qualitative signatures of confidence have limited applicability, what is the way forward? One option available to confidence researchers is to discover more signatures, being careful to find the specific conditions under which they are expected. Confidence experimentalists should then make sure to look for such signatures only when their tasks satisfy the specified conditions (e.g., stimulus distribution type, noise level). However, for researchers interested in testing the BCH, we do not necessarily advocate for this course of action because, even when applied to relevant experiments, the presence or absence of qualitative signatures provides an uncertain amount of evidence for or against the BCH. Testing for the presence of qualitative signatures is a weak substitute for accumulating probabilistic evidence, something that careful^32^ quantitative model comparison does more objectively. Testing for signatures requires the experimenter to make two subjective judgments. First, the experimenter must determine whether the signature is present, a task potentially made difficult by the fact that real data is noisy. Second, the experimenter must determine how much evidence that provides in favor of the BCH, and whether further investigation is warranted. By contrast, model comparison provides a principled quantity (namely, a log likelihood) in favor of the BCH over some other model^1,2,7^. Given the caveats associated with qualitative signatures, it may be that, as a field, we have no choice but to rely on formal model comparison.

## APPENDIX A: Derivation of Bayesian confidence

As described in Section 2, if an observer’s confidence behavior is Bayesian, it is a function of the posterior probability of the most probable category. By Bayes’ rule,

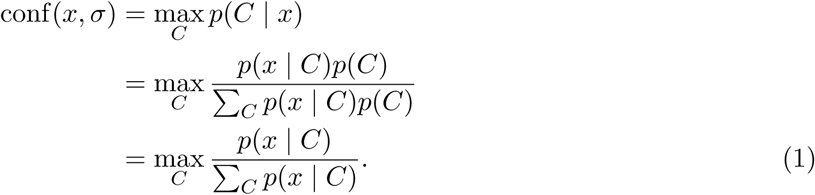

In the last step, we eliminated the prior because each category is equally likely (i.e., *p*(*C* = 1) = *p*(*C* = −1)) and we assume that the observer knows this. We now derive the task-specific likelihood functions *p*(*x* | *C*) used in our simulations. The observer does not know the true stimulus value s, but does know that the measurement is drawn from a Gaussian distribution with a mean of *s* and s.d. *σ*. Using this knowledge, the optimal observer marginalizes over *s* by convolving the stimulus distributions with their noise distribution:

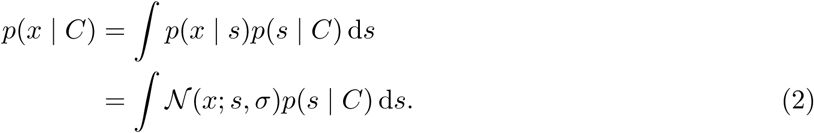

For uniform category distributions, we plug 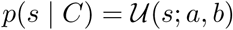 into Equation (2) and simplify:

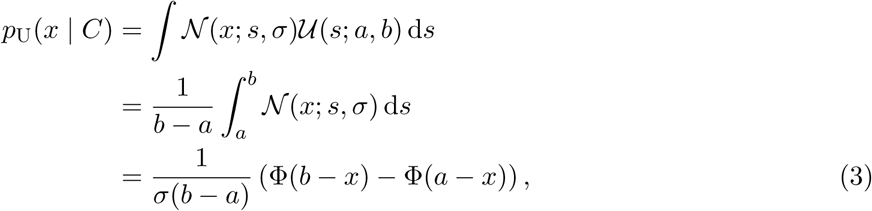

where Φ is the cumulative distribution function of the standard normal distribution. For Gaussian category distributions, we plug 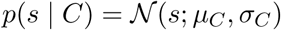 into Equation (2) and simplify:

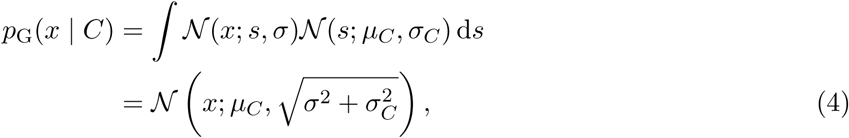

using *σ_C_* = 0 if stimuli from a given category always take on the same value *μ_C_*.

Finally, plug the task-appropriate likelihood function (Equation (3) or Equation (4)) into Equation (1).

## APPENDIX B: Notation table

Because some of our notation relates to that used in Hangya et al.^14^, we provide this table to enable easier comparison between the two papers. In some cases, the variables are not exactly identical: the terms in Hangya et al. may be more general. This does not affect the validity of our claims. For consistency, we always describe their work using our notation.

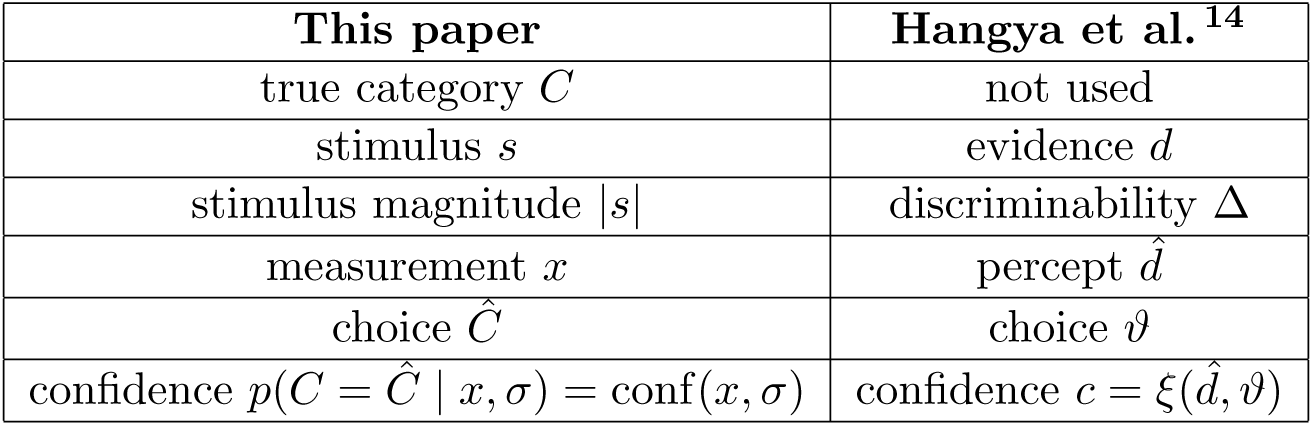

## APPENDIX C: Simpler proof of Hangya et al.^14^ lemma

The proof of the 0.75 signature depends on a lemma proved by Hangya et al.^14^: *Integrating the product of the probability density function f and the distribution function F of any probability distribution symmetric to zero over the positive half-line results in 3/8*:

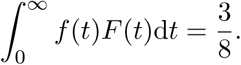

There is a shorter proof of the lemma, which is as follows. Use integration by parts, and that *f* (*t*) = *F*′(*t*) by definition:

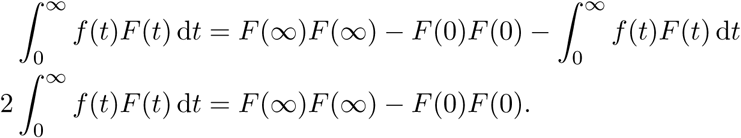

Because *F* is a cumulative distribution function of a probability distribution symmetric across zero, *F*(∞) = 1 and 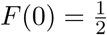:

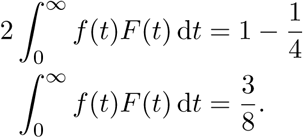

## ACKNOWLEDGEMENTS

The authors would like to thank Luigi Acerbi, Rachel N. Denison, Andra Mihali, and Joaquin Navajas for helpful conversations and comments on the manuscript; Bas van Opheusden for the simple proof of the lemma; and Rachel Adler for some clever real-life examples of overlapping categories. This material is based upon work supported by the National Science Foundation Graduate Research Fellowship under Grant No. DGE-1342536.

## CODE AVAILABILITY

All code used for simulation and plotting is available at GitHub/wtadler/confidence/signatures.

## Footnotes

I Restating this logic in probabilistic terms: A signature being a necessary condition for the BCH implies that *p*(signature observed | BCH is true) = 1. A signature being an insufficient condition implies that *p*(signature observed | BCH is false) > 0. By Bayes’ rule, for signatures that are both necessary and insufficient, *p*(BCH is true | signature(s) observed) will increase with the observation of each signature but will never reach 1.

II Note that our assumption that confidence and category choice are deterministic functions of *x* amounts to an assumption that there is no noise at the action (i.e., reporting) stage.

III Kepecs and Mainen^17^, Insabato et al.^16^, and Fleming and Daw^11^ call it the (folded) “X-pattern.”

IV Our finding is distinct from that of Insabato et al.^16^, who show that the divergence signature would not be predicted under a non-Bayesian model in which the observer uses two measurements on each trial. Our analyses only concern Bayesian models in which the observer has a single measurement on each trial. Our finding is also distinct from that of Fleming and Daw^11^, who show that the divergence signature would not be predicted if the experimenter could plot confidence as a function of the internal measurement *x*. Our analyses only concern confidence as a function of stimulus magnitude |*s*| which, unlike *x*, is known by the experimenter.

V Their original assumption actually reads, “for any given confidence *c*, the relative frequency of percepts mapping to *c* by *ξ* changes monotonically with evidence discriminability for any fixed choice.” In our terminology, this is equivalent to saying that, as |*s*| increases, the frequency of reporting any particular level of confidence changes monotonically. This is not correct even in the case of uniform stimulus distributions; for example, at low noise, as discriminability increases, the frequency of medium-confidence reports will increase and then decrease. Their restatement of this assumption specifically for incorrect choices, which we cite in the main text, is correct for non-overlapping stimulus distributions. Because they restate the assumption correctly, their following argument holds except under the scenario described in the main text.

VI In the Bayesian model, observers use their knowledge of their uncertainty. In this alternative standard signal detection theoretical model^13,17^, observers ignore uncertainty, making confidence only a function of the distance between the measurement and the decision bound. Previous studies have referred to similar models as Fixed^1,7,35^ or Difference^3^.

VII However, because the orientations were drawn such that the mean orientation of each set was the same for all trials in a category, there was no variance over the stimulus variable of interest (the per-trial mean) within categories. Therefore, what they describe as stimulus variance factors into a Bayesian model of confidence (and into their non-Bayesian decision model) only by changing measurement noise. Additionally, because there is no variance over stimulus magnitude within categories, they are unable to determine whether divergence signature #1 is present in their data.

VIII This corresponds to Navajas et al.^29^, as desciibed in Section 5.

IX In their model, which they label “normative,” the observer continually updates a weighted average of each stimulus with the previous average. This model is not equivalent to (nor a supermodel of) the optimal model, which keeps a running sum of stimuli, dividing by *N* for each stimulus or at the end of the trial. They motivate their non-Bayesian model by the observation that recent samples have a relatively higher influence on subject decisions, but do not show fits of a fully Bayesian model to their data.

X Kiani et al.^20^ also note the lack of the divergence signature in their data, but because their stimuli have variable duration, optimality is more complicated to characterize^9^, and the explanation we offer here may not apply.

